# Exploration of interaction scoring criteria in the CANDO platform

**DOI:** 10.1101/591578

**Authors:** Zackary Falls, William Mangione, James Schuler, Ram Samudrala

## Abstract

**Background:** Drug discovery is an arduous process that requires many years and billions of dollars before approval for patient use. However, there are a number of drugs and human ingestibles approved for a variety of indications/diseases that can be potentially repurposed as new treatments for others, decreasing the time and cost required.

**Methods:** CANDO (Computational Analysis of Novel Drug Opportunities) is a platform for shotgun, multitarget drug discovery and repurposing. The CANDO platform scores interactions between 46,784 proteins structures and 3,733 human use compounds using a bioinformatic docking protocol to generate compound-proteome interaction signatures that are then compared to identify candidates for repurposing. Benchmarking of the platform is accomplished by comparing the compound-proteome interaction signatures and determining whether signatures corresponding to pairs of drugs approved for the same indication fall within particular cutoffs.

**Results:** We have altered the scoring function of bioinformatic docking protocol in the newest version of our platform (v1.5) to use the best OBscore for each compound-protein interaction, resulting in an increased benchmarking accuracy from 11.7% in v1 to 12.8% in v1.5 for the top10 cutoff, the most stringent one used, and correspondingly from 24.9% to 31.2% for the top100 cutoff.

**Conclusions:** The change in the interaction scoring and other bug fixes in CANDO v1.5 have resulted in improved benchmarking performance, making the platform more effective at predicting novel, therapeutic drug-indication pairs.

## Background

Drug discovery is an arduous process that requires many years of effort and costs billions of dollars before new ones are approved for patient use.^1;2^ Recent data indicate that the average cost and time to market for a new drug are about $3 billion and 14 years, respectively.^3;4^ New paradigms are therefore imperative to make drug discovery more efficient and financially sustainable.

As of 2013, there were ≈ 1,453 human use drugs FDA approved for a variety of indications/diseases with an accompanying trove of data on their safety profiles and efficacy.^5^ A vast majority of these drugs are small molecules, which are inherently promiscuous in their potential interactions with macromolecules in their environment, resulting in undesirable off-target or side effects.^6–11^ The multitargeting nature of small molecules, and the presence of these off-target effects, provides support for the concept of repurposing drugs for indications they are not approved for.^7;11–15^ The cost, time, and most importantly, risk, to go from “bench to bedside” for such repurposed drugs are significantly decreased.

Drug repurposing methodologies can be categorized into two classes: experimental and ontological.^16^ Experimental methodologies include direct investigation via *in vitro* and *in vivo* screening, as well as hypothesis driven computational approaches such as molecular modelling.^17–22^ Ontological methodologies leverage growth in databases and advances in natural language processing, using known information about drugs (such as indication associations, biological targets, etc.) to deduce previously unknown relationships between the drugs and other indications.^16;23–28^ Our Computational Analysis of Novel Drug Opportunities (CANDO) platform is a hybrid methodology that uses a combination of knowledge-based approaches (known associations between drugs and indications, ^29^ potentials of mean force,^30^ and sequence and structure databases^31^) as well as protein structure and interaction modelling^32–34^ and ligand similarity comparisons ^35^ to predict novel relationships between pairs of compounds and indications.^7;14;15;36–38^

CANDO is a platform for multitarget shotgun drug repurposing.^7;11;14;15;36–39^ The first version (v1) of the CANDO platform implemented a modelling pipeline to predict interactions between 46,784 protein structures and 3,733 human use compounds. Various protocols, representing software components, are implemented within each pipeline to calculate an interaction score for each drug-protein pair corresponding to the potential binding affinty. Applied across entire proteomes, this results in compound-proteome interaction signatures that are then compared and ranked according to similarity. We then generate new indication associations for drugs based on the similarity of their interaction signatures to drugs with a known indication, *i.e.*, make predictions about putative repurposeable therapeutics for every indication with at least one known drug. Furthermore, we quantify the expected accuracy of our predictions by performing a leave-one-out benchmarking procedure which determines whether an associated drug for each known drug-indication pair is captured within a cutoff of a list of compounds sorted by proteomic signature similarity to the “left out” drug.

In the v1 platform, we used an interaction scoring protocol that integrated bioinformatics and cheminformatics tools to calculate ≈ one billion scores. We updated our platform to v1.5 by exploring the use of different bio- and cheminformatic software to vary these interaction scores to discover the best performing scoring protocol. The pipelines implementing these new scoring protocols were subsequently benchmarked, the results of which are reported here.

All of the pipelines with the new interaction scoring protocols in CANDO v1.5 yield promising benchmark performance. However, there is some variance depending on how many top putative drug candidates are generated and benchmarked: At the lowest cutoff (top10 putative drug candidates), the pipeline with the best performance uses only the cheminformatic interaction score. At higher cutoffs (top25 - top100), the pipeline with the best performance combines the bioinformatic and cheminformatic outputs for the interaction scores. These results help guide future experimental validation studies of the platform by enabling us to select the appropriate interaction scoring protocol based on the number of putative drug candidates to be tested.

## Methods

### Putative drug library creation and indication mapping

A virtual library of FDA approved and other human use compounds was created using multiple sources including DrugBank,^40^ NCGC Pharmaceutical Collection (NPC),^41^ Wikipedia, and PubChem.^42^ Each molecule is converted to a 3D structure using MarvinBeans molconverter v5.11.3 from ChemAxon.^43^ Cactvs Chemoinformatics Toolkit ^44;45^ from Xemistry was used to generate unique InChIKeys (International Chemical Identifier Keys) from the clean 3D chemical structures and each compound InChIKey was compared to all others to eliminate redundancy in the drug library. The final library contains 3,733 human use compounds, including 1387 approved drugs that map to 2,030 diseases/indications. Drug indication associations were obtained from the Comparative Toxicogenomics database.^29^ 1,439 indications (out of the 2,030) are associated with two or more drugs and are used to perform the leave-one-out benchmarking of the platform described below.

### Protein structure library creation and binding site predictions

A total of 46,784 solved and modeled protein structures comprise the library of potential macromolecular targets in the CANDO platform. The solved protein structures (31,145) are obtained from the Protein Data Bank (PDB)^31^ and the modeled structures (15,639) are generated using the I-TASSER software^46^. The bioinformatic tool COFACTOR^32–34^ is used to assess binding site similarity (Figure 1A) by comparing the protein of interest to a library of template proteins from the PDB to determine the best match for potential binding sites.^47^ The metric that defines the local structure and sequence similarity between the target and template binding sites is the binding site similarity score (BSscore).^34^

**Figure 1:**
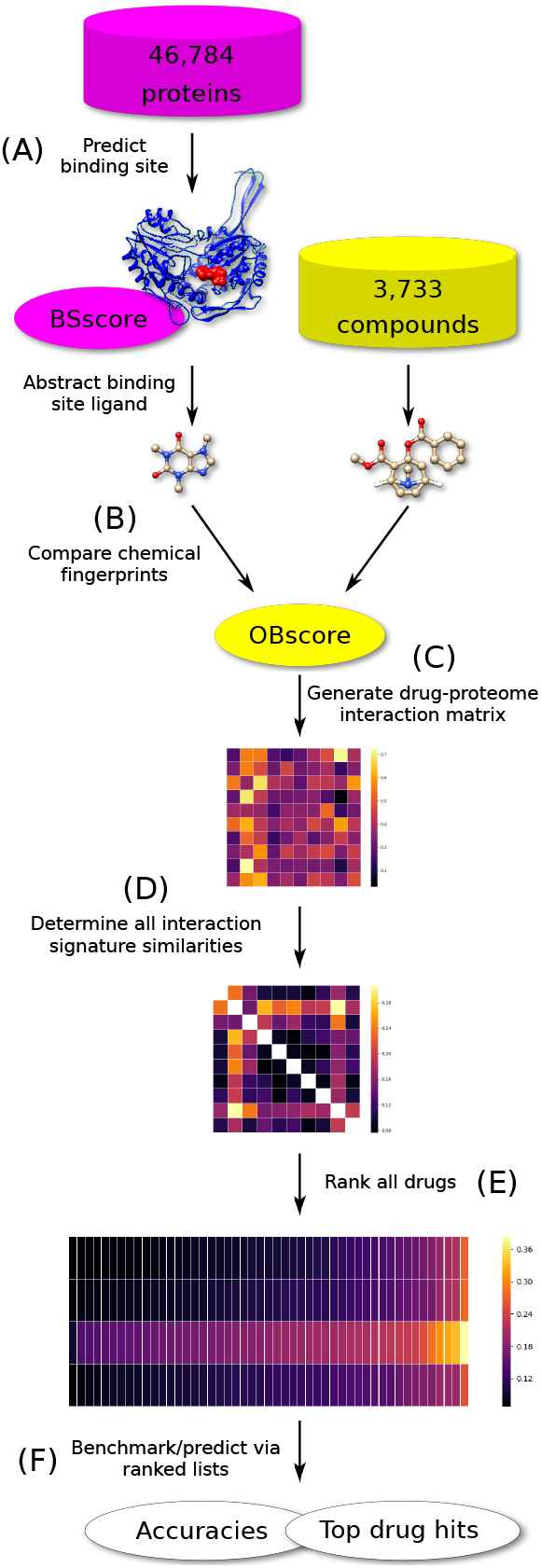
CANDO platform workflow. (A) Binding sites are predicted for each of the 46,784 proteins in the CANDO protein structure library, resulting in a BSscore. (B) The native ligand in the predicted binding site is compared to all 3,733 compounds in the CANDO putative drug library by calculating the chemical fingerprints for each structure, resulting in an OBscore. (C) Each compound-protein interaction is given a score based upon the OBscore and/or BSscore, which is then used to populate the interaction matrix. (D) The similarity score between every pair of compound-proteome interaction signatures (the vectors of 46,784 interaction scores) is calculated by root-mean-squared deviation (RMSD) which are then used to populate the compound-compound similarity matrix. (E) The compound-compound similarities are sorted and ranked by RMSD. (F) Benchmarking is accomplished by measuring the recovery rate of the known approved drugs, *i.e.*, per indication accuracies are obtained based on whether or not pairs of drugs associated with the same indication can be captured within a certain cutoff of each of their ranked compound similarity lists; other similar compounds that fall within a particular cutoff are hypothesized to be repurposeable drugs and serve as predictions. The CANDO platform utilizes a proteomic approach for drug repurposing, with the hypothesis that drugs with similar interaction signatures will behave similarly.

### Chemical fingerprint identification and comparison

The FP4 fingerprinting method in Open Babel is used to determine the cheminformatic structural similarity score (OBscore)^35^ between all binding site ligands from COFACTOR and structures in the putative drug library (Figure 1B). We chose Open Babel to determine structural fingerprints in CANDO v1.5^48^ as it is an established suite of open source software.^35^

### Generation of compound-proteome interaction signatures

We quantify the compound-protein interaction strength using combinations of the OBscore and/or BSscore described above. When applied to the corresponding libraries, this generates a compound-protein interaction matrix (Figure 1C), where each row of this matrix, *i.e.* the compound-proteome interaction signature, describes how each compound interacts with the entire multiorganism protein library.

In CANDO v1.5, we use the OBscore and BScore to populate the interaction matrix for the following pipelines: Best OB, Best BS, Best OB+BS, and Best OBxBS. The values in the matrix for each compound-protein interaction in the first two pipelines use the OBscore; Best OB is the highest OBscore between the compound and all predicted binding site ligands for each protein, while Best BS is the OBscore that corresponds to the best local binding site prediction using COFACTOR. The last two pipelines involve adding and multiplying the OBscore and BSscore for each compound-protein interaction; the highest sum or product between the compound and the predicted binding site ligands was chosen as the interaction score.

### Calculating interaction signature similarities

The similarity between every compound-proteome interaction signature is compared to all other signatures (Figure 1D) using the root-mean-square deviation (RMSD). This procedure generates a symmetric matrix (with zeroes along the diagonal) of similarity scores that are hypothesised to represent how functionally similar each compound is to all the others in the context of the protein structure library.

### Ranking drug lists and benchmarking metrics

The RMSDs in each row of the compound-compound similarity matrix are sorted to yield ranked similarity lists for each compound (Figure 1E). Each drug associated with an indication is left/held out and checked to see if it is captured within a certain cutoff in the ranked list to any of the other remaining ones [associated with that indication] (Figure 1F). The cutoffs used typically are top10, top25, top50, and top100, reflecting the top ranked 10-100 similar compounds for a given drug.

This procedure is repeated iteratively for all drugs associated with every indication for a particular cutoff, resulting in the indication accuracy. Mathematically, indication accuracy is calculated using the formula 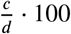, where *c* is the number of times at least one drug with the same indication was captured within a particular cutoff and *d* is the total number of drugs approved for that indication. Taking the mean of these accuracies (for all 1439 indications with at least two approved drugs) gives the average indication accuracy for a pipeline at a particular cutoff.

The other benchmarking metrics used are the average pairwise accuracy which is a weighted average of all indication accuracies based upon the number of approved drugs for each indication, and indication coverage, which is the number of indications with a non-zero accuracy (i.e., at least one approved drug that was left out was successfully recaptured within a cutoff).

### Differences between versions 1 and v1.5 of the CANDO platform

For v1.5, we use Open Babel for the chemical fingerprint comparison between all compounds and predicted binding site ligands for each protein, compared to using OpenEye ROCS in v1.^36^ Pipeline modifications have been made to leverage OBscore and/or BSscore to populate the interaction matrix in multiple pipelines for CANDO v1.5, whereas only the BSscore was used in CANDO v1 to calculate compound-protein interactions.

By removing the cutoffs for interaction scores (BSscore and ROCSscore in CANDO v1^36^), we decreased the number of compound-protein interactions with zero scores, which we empirically determined had a negative effect on benchmarking performance (Figure 2). Additional minor modifications have been made in CANDO v1.5 software to reduce the number of compounds with all-zero proteome interaction signatures.

**Figure 2:**
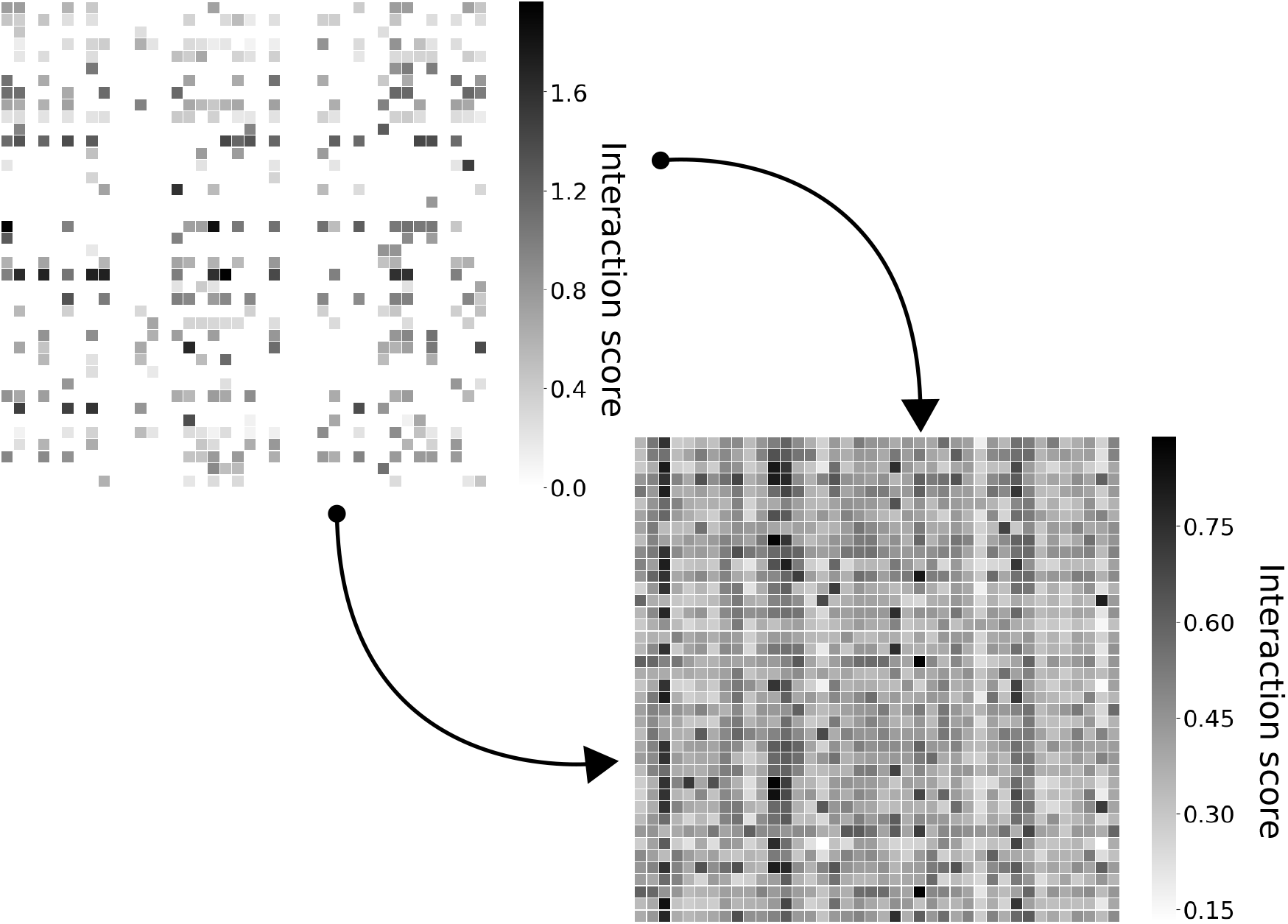
Increased signal in v1.5 interaction matrix. In v1, about 800 compounds had all-zero signatures, wheres in v1.5 all of these compounds have non-zero signatures. The average indication accuracy increased 1.1% (from 11.7% to 12.8%) for the top10 cutoff, and by 6.3% for the top100 cutoff (from 24.9% to 31.2%). Upgrading the CANDO v1 pipeline to v1.5 has resulted in the signal-to-noise ratio being increased (i.e., more interactions calculated) by about 20% for the compound-proteome interaction matrix.

### Generation of random control interaction matrices

We generated random compound-proteome interaction matrices to compare the efficacy of v1 and the v1.5 pipelines against a control. For each compound-protein interaction score we randomly selected a value from a uniform distribution between 0.0 and 1.0 to populate a 3,733 (compounds) by 46,784 (proteins) interaction matrix. We benchmarked this matrix, as discussed in a previous section, to ascertain the benchmarking metrics (average indication accuracy, average pairwise accuracy, and indication coverage) for all cutoffs (top10, top25, top50, and top100). This protocol was repeated 100 times and the resulting averaged metrics were used as the random control.

We calculated the hypergeometric distribution for our leave-one-out benchmark at each cutoff as an additional control using Eq. 1.

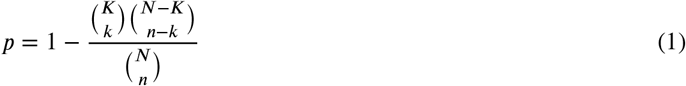

where *p* is the probability of recapturing at least one drug approved (*k*) for the same indication as the “left out” drug in the top*n* cutoff, considering a population (*N*) of 3732 compounds and *K* being the number of approved drugs for the indication. The probability was calculated and averaged across all indications, for which the number of approved drugs, *K*, varies.

## Results and discussion

We generated compound-proteome interaction matrices using the BSscore and OBscore interaction scoring schemes (Section) to implement the following pipelines: Best OB, Best BS, Best OB+BS, and Best OBxBS. These pipelines were compared to the one used in CANDO v1, as well as random controls, with respect to benchmarking performance using three evaluation metrics: average indication accuracy, average pairwise accuracy, and indication coverage.

### Increasing the signal in the matrices and pipeline in CANDO v1.5

Using the v1.5 pipeline we decreased the noise in the compound-proteome interaction matrix relative to v1. Specifically, in v1 more than 800 compounds in our library had completely null interaction signatures (as shown in Figure 2), i.e., every compound-protein interaction for the ≈ 800 compounds received a score of 0.0. We reduced that number to less than 50 compounds, all of which are chemical ions that fail in the current version of the bioinformatic docking protocol. In addition, ≈ 66% of all compound-protein interaction scores in v1 was assigned a score of 0: Each compound-proteome interaction signature in v1, on average, contained 38,492/46,784 (median = 36,778) null or zero interaction scores, whereas the average number of null interaction scores within v1.5 has decreased significantly to 792 (median = 40).

These changes resulted in an increase from 11.7% in v1 to 12.8% in v1.5 for the top10 average indication accuracy, corresponding to 9.4% increase between versions. Furthermore, a greater improvement in accuracy between versions was observed at higher cutoffs, with a 25% increase for the top100 average indication accuracy. This indicates a greater capability of our platform to recapture known drugs for each indication, as well as reflects the increased capability of predicting putative repurposable drugs for all indications.

### Variation of OBscore and BSscore threshold values

In v1, we used a threshold value of 1.1 for the ROCSscore and BSscore to determine if a protein-compound interaction would occur, based on an analysis of structure-ligand complexes.^36^ For v1.5, we benchmarked the Best OB matrix for each incremental increase in OBscore (0 to 1) and BSscore (0 to 2) values individually to determine how these thresholds influence the overall benchmarking result.

As shown in Figure 3, both OBscore and BSscore thresholds were increased incrementally and independently from 0 to their corresponding theoretical maxima of 1 and 2 respectively. As the OBscore threshold is increased, the resulting accuracies and coverages decrease, eventually approaching zero (Fig. 3). This is because the ligand and compound must have near identical chemical similarity to have a high OBscore. However, few of the compound in the CANDO library are chemically similar to the ligands present in the binding sites of the template proteins from COFACTOR, let alone identical (where the OBscore would be 1). The results from Figure 4 indicates that as the OBscore threshold increases, the number of non-zero OBscores in the interaction matrix will decrease, and the average indication accuracies and coverage will decrease correspondingly (Figure 3).

**Figure 3:**
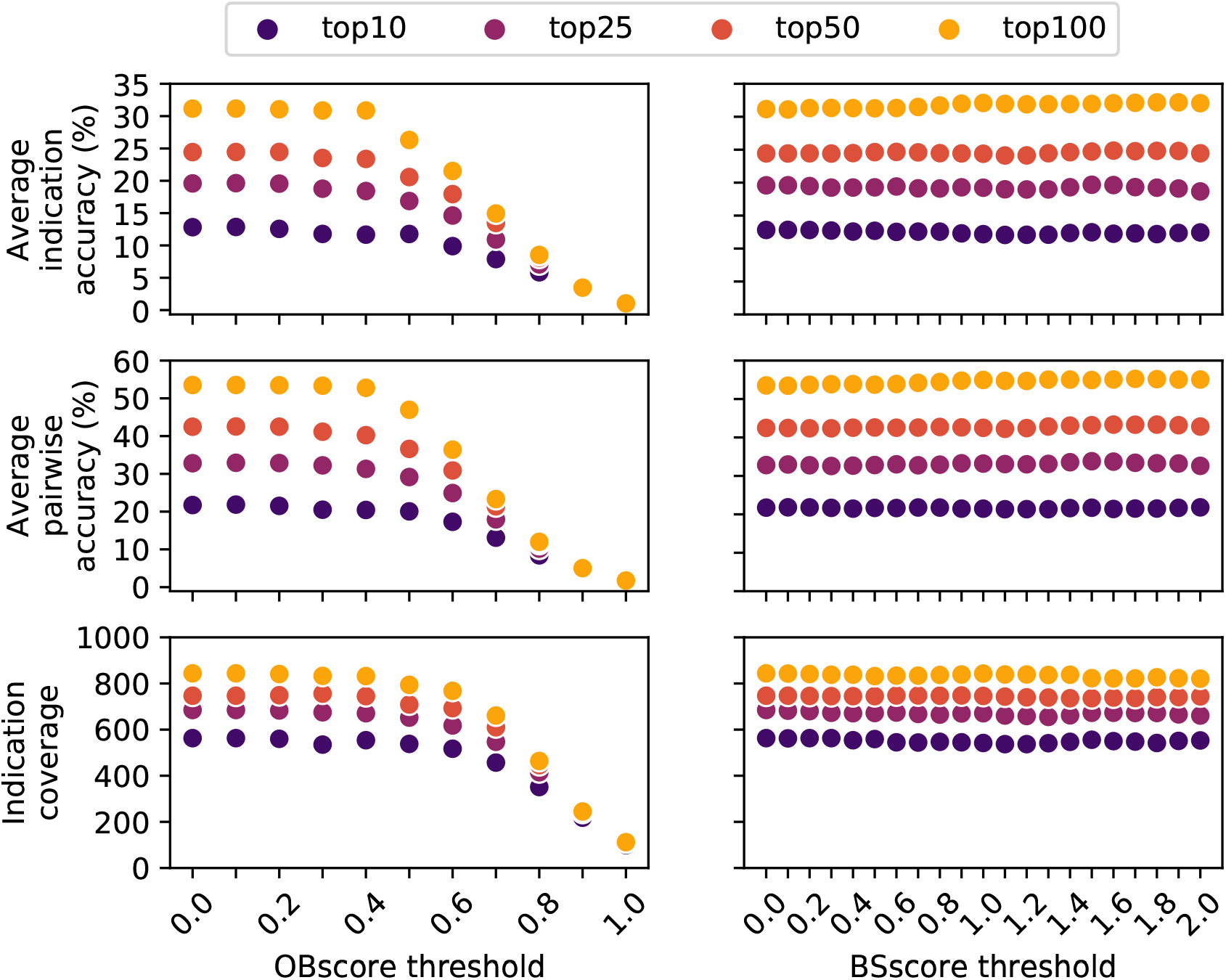
Change in benchmarking performance when varying the OBscore or BSscore thresholds. The OBscore and BSscore thresholds were set to a value between 0.0-1.0 and 0.0-2.0, respectively; scores above the threshold value were used to populate the compound-proteome interaction matrix and scores below were set to zero. Matrix generation and benchmarking was performed over the full range at increments of 0.1 for OBscore and 0.2 for BSscore. The average indication accuracy, average pairwise accuracy, and indication coverage are shown for four compound-compound similarity list cutoffs: top10 (purple), top25 (magenta), top50 (red), and top100 (yellow). The plots for the OBscore show that as the threshold is increased towards the theoretical maximum of 1.0, the resulting accuracies diminish to nearly 0%. This is because as the threshold becomes more stringent, the number of compounds that have a strong similarity (OBscore greater than the threshold) to the binding site ligands approaches zero, therefore more of the drugs will have near–zero proteome signatures. If there is no signal to discern compound–proteome interaction signature similarity, then the benchmarking produces decreasing accuracies with each increment to the threshold value. In contrast, the BSscore plots show that as the threshold is increased toward the theoretical maximum of 2.0, there is negligible fluctuation in the average indication and pairwise accuracies because there exist predicted binding sites for most proteins in our library that receive a BSscore of 2.0 from COFACTOR. Based upon these results, we used a lower cutoff for the OB and BSscores to obtain the best benchmarking performance.

In contrast, the incremental increase in the BSscore threshold does not significantly affect benchmarking performance (Fig. 3). A large portion of the protein structure library used by the CANDO platform consists of solved PDB structures that overlap highly with the COFACTOR template library. This results in a considerable amount of signal remaining at the highest thresholds (Figure 4). However, the indication coverage appears to decrease by ≈ 30 as the threshold is increased, meaning the high OBscore cutoff is resulting in the average indication accuracies to remain consistent, however, the accuracy is averaged over fewer number of indications [this sentence is unclear and needs to be rewritten].

**Figure 4:**
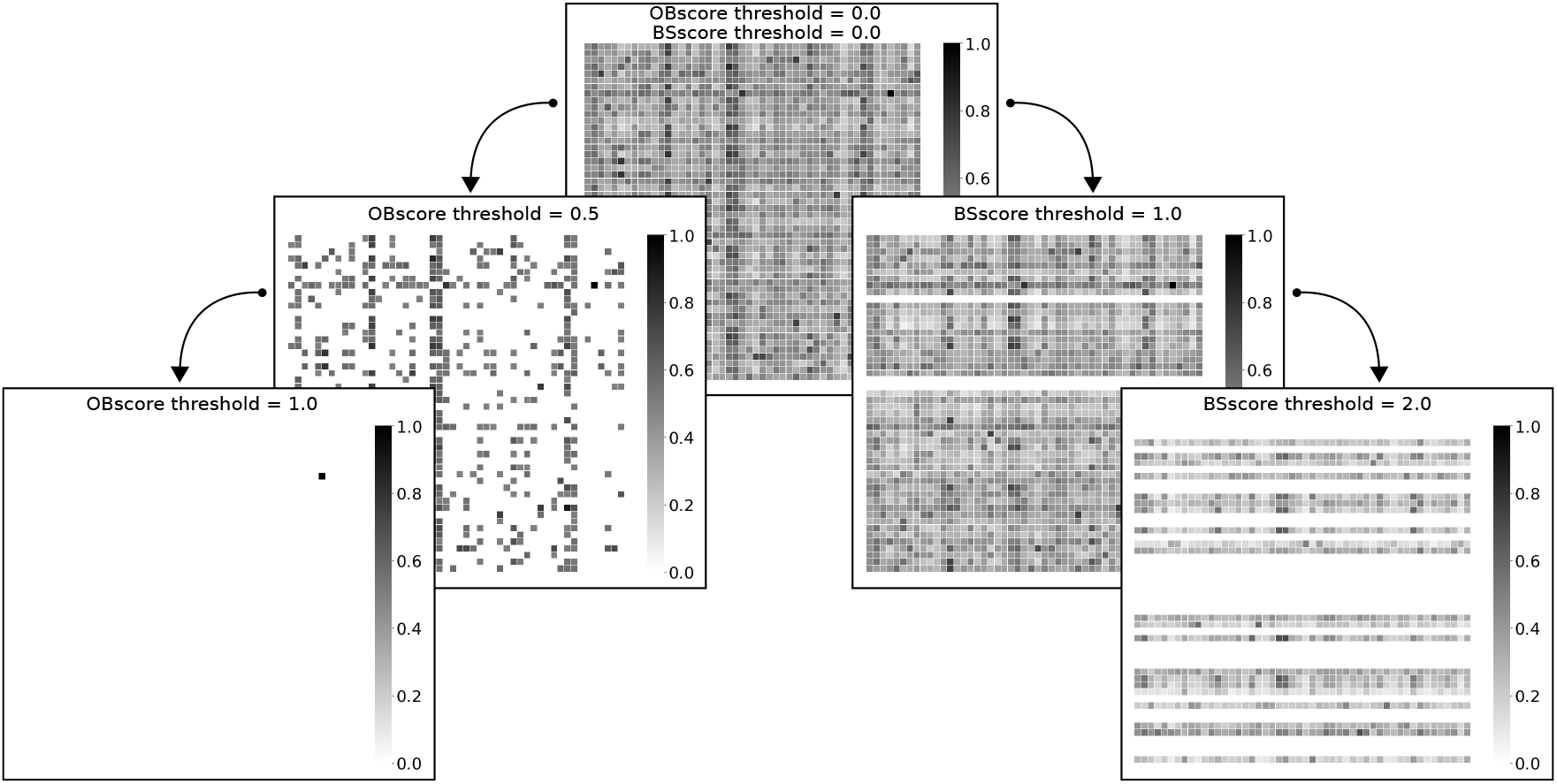
Increase in signal in the interaction matrices as a function of score thresholds. A small section of the matrix that corresponds to no threshold for either BSscore or OBscore is shown top centre. A two step threshold increase of OBscore (0.0 → 0.5 → 1.0) is shown to its left and a two step threshold increase of the BSscore (0.0 → 1.0 → 2.0) is shown to its right. As the OBscore threshold is increased, the resulting matrix contains many null values (interaction score of zero) leading to diminished accuracies (Figure 3. In contrast, increasing the BSscore threshold yields a sufficient amount of signal, with the corresponding benchmarking performance largely unchanged. Overall, these results indicate that using lower BSscore and OBscore thresholds yield better benchmarking performance.

### Comparison of random control and hypergeometric distribution accuracies

The average accuracies from the hypergeometric distributions for the top cutoffs (top10, top25, top50, top100) are 2.0, 5.1, 9.7, and 17.3%, respectively. The average indication accuracies resulting from the randomly generated compound-proteome interaction matrices converge to the same value as the hypergeometric distribution accuracies. This result verifies the benchmarking protocol and confirms the other metrics for determining accuracy (average pairwise accuracy and indication coverage) are convergent for the randomly generated results.

### Comparison of v1 and Best OB pipelines

The CANDO v1.5 Best OB pipeline average indication accuracy is higher at all cutoffs when comparing to the pipeline from CANDO v1, increasing from 11.7% to 12.8% for the top10 cutoff. The relative increase in average indication accuracy for the remaining cutoffs are 3.0% (top25), 4.1% (top50), and 6.3% (top100). The indication coverage for Best OB is greater than v1 at all cutoffs (30 – 70 more non-zero indications) except top10, where the coverage for v1 and Best OB is about the same at 562 and 563 indications, respectively.

We calculated the Kolmogorov–Smirnoff test p-value to determine that the distribution of indication accuracies was significantly different between v1 and Best OB pipelines for all cutoffs (Figure 5). Furthermore, the distributions in Figure 5 show that the accuracies for Best OB, relative to v1, are skewed to the right, i.e., Best OB has a greater number of indications with accuracies > 50%.

**Figure 5:**
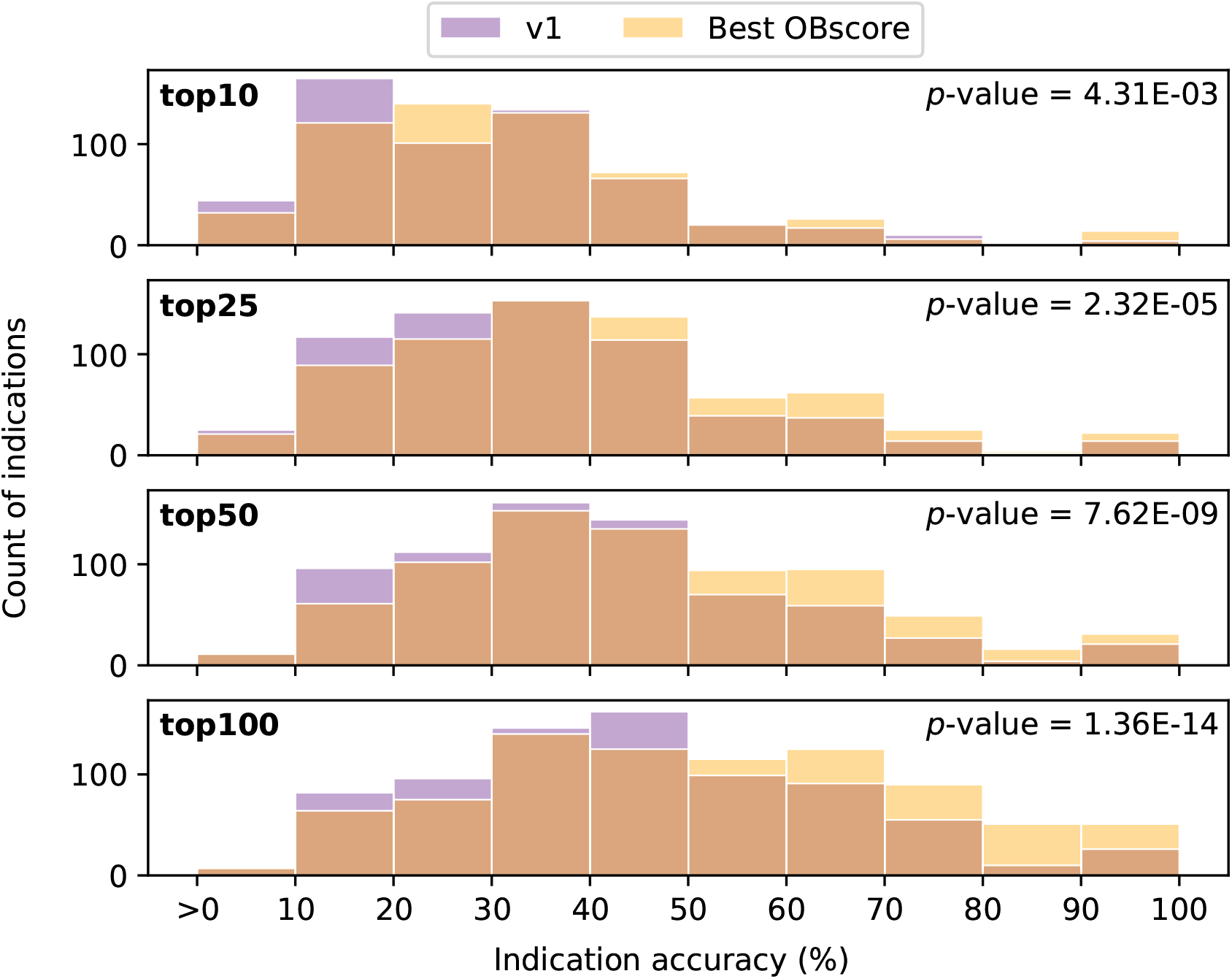
Comparison of indication accuracy distributions. A histogram of the non-zero accuracy values for the v1 (blue) and Best OBscore (yellow) pipelines at four cutoffs is plotted. The Kolmogorov-Smirnoff test, used to determine similarity (or lack thereof) of two distributions, indicates that the two pipelines have significantly different distributions of indication accuracies (p-value < 0.05). The newer v1.5 Best OB pipeline outperforms its predecessor, yielding a greater number of indications with accuracies *>* 50%.

### Comparison of all pipelines

Figure 6 shows the accuracies and coverages of all five pipelines at different cutoffs. All scoring metrics in v1.5 did comparably well to one another and better than the pipeline used in the CANDO v1 platform. Best OB produces the highest average indication accuracy of 12.8% and 19.6% for the top10 and top25 cutoffs. At higher cutoffs, the Best BS, OB+BS, and OBxBS pipelines perform better for the average and pairwise indication accuracy metrics, with OBxBS having the highest average indication accuracy of 31.8% at the top100 cutoff.

**Figure 6:**
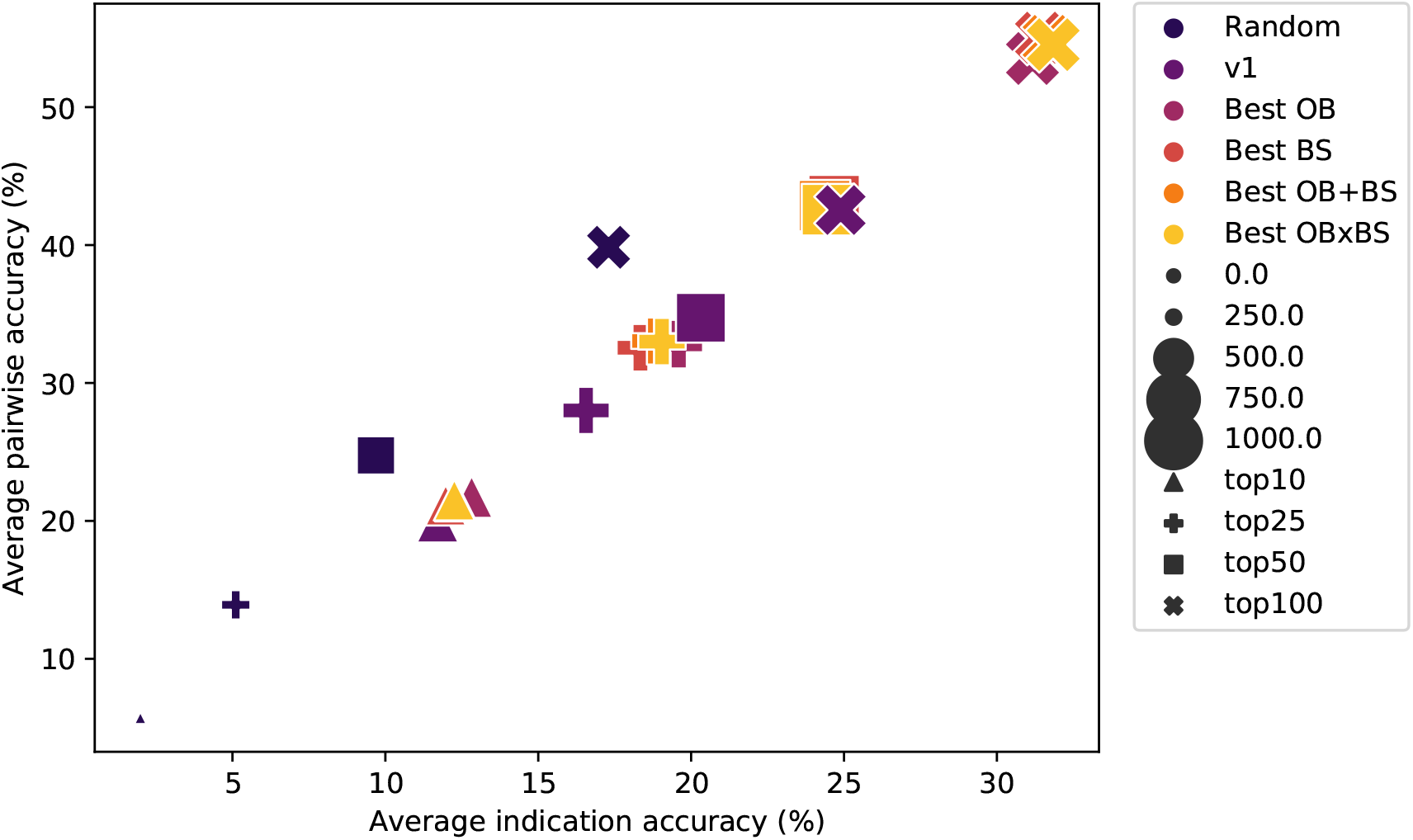
Accuracies and coverages of five CANDO pipelines at various cutoffs using different compound-protein interaction scoring metrics. Random (navy) is a random control that is calculated using the average of 100 randomly generated interaction matrices. v1 (purple) is the pipeline from the first version of the CANDO platform which used the BScore to quantify compound–protein interactions, with the chemical similarity comparison done using OpenEye ROCS.^7;36^ The Best OB pipeline (magenta) uses the highest OBscore for each compound-protein interaction. Best BS (red) uses the OBscore corresponding to the fingerprint comparison between each compound and highest BSscore binding site ligand for each protein. Best OB+BS (orange) is the highest summation of OBscore and BSscore for each compound-protein interaction, and Best OBxBS (yellow) is the highest product. The average indication and pairwise accuracies for the top10 (triangle), top25 (cross), top50 (square), and top100 (X) cutoffs are shown for each pipeline. The size of the cutoff symbol corresponds to the indication coverage. By varying the interaction scoring scheme in v1.5, we are able to discern that the Best OB protocol results in the highest benchmarking performance, particularly at the top10 and top 25 cutoffs.

The Best OB pipeline average indication accuracy is five times greater at the top10 cutoff when compared to the random control (2.0 to 12.8%). This trend remains consistent as the cutoff increases, with relative differences between random control and Best OB of 14.5, 14.7, and 13.9% for the top25, top50, and top100 cutoffs. The average pairwise accuracies and indication coverage are also higher for the Best OB pipeline when compared to the random control with a pairwise accuracy increase from 5.7 to 21.7% and coverage increase from 238 to 563 at the top10 cutoff. The relative increases between the Best OB and random control are 18.9, 17.8, and 13.7% for average pairwise accuracy and 305, 245, and 207 for the indication coverage at the remaining three cutoffs.

## Conclusions

The CANDO platform is used to generate top ranking putative drug candidates for every indication. These candidates need to be experimentally validated to ensure they represent potential leads and eventually repurposed drugs for a specific indication. Our results suggest that for preclinical validations of 25 or fewer compounds, the Best OB pipeline, which has the highest average indication accuracy, pairwise accuracy, and indication coverage at the top10 and top25 cutoffs, should be used to generate putative drug candidates. In contrast, the results show that at higher cutoffs (top50 and top100) the Best OBxBS and Best OB+BS pipelines yield better benchmarking performance, indicating that these two pipelines should be utilized for validation studies consisting of 50 or more putative drug candidates.

The update from v1 to v1.5 has led to improved benchmarking performance in the CANDO platform by increasing the signal-to-noise ratio of the interaction matrix resulting in a significant increase in the average indication accuracy. Furthermore, varying the OBscore and BSscore thresholds during the interaction matrix generation showed that using any thresholds for these scores results in the reduced signal which is reflected in benchmarking performance.

Other possible scoring protocols need to be explored to determine if OBscore and BSscore most accurately quantify the compound-protein interactions. Further studies with different cheminformatic and bioinformatic tools may also provide further insight into the behaviour of the platform and are currently underway, which demonstrate that continued development of CANDO by adding novel features and pipelines greatly increases its predictive power for future drug repurposing efforts particularly when these other pipelines and optimization techniques are used in combination.^11;39^

Overall, our results illustrate the improved benchmarking performance of the updated CANDO v1.5 platform and its structure-based pipelines relative to v1, which in turn translates to greater predictive power for shotgun drug repurposing and mechanistic understanding. The top putative drug candidates and targets generated by these newer pipelines in v1.5 will aid us in discovering novel treatments and mechanisms for specific indications in future validation studies.

## Declarations

**Abbreviations**
CANDO: Computational Analysis of Novel Drug Opportunities
OBscore: Open babel score
BSscore: Binding site score
PDB: Protein Data Bank
RMSD: Root mean squared deviation

### Ethics approval and consent to participate

Not applicable

### Consent for publication

Not applicable

## Availability of data and materials

The datasets supporting the conclusions of this article are available in the CANDO repository, http://protinfo.org/cando/results/v1_5.

### Funding

This work has been supported by a National Institute of Health Director’s Pioneer Award (DP1OD006779), a National Institute ofHealth Clinical and Translational Sciences Award (UL1TR001412), a National Library of Medicine T15 Award (T15LM012495), a National Cancer Institute/Veterans Affairs Big Data-Scientist Training Enhancement Program Fellowship in Big Data Sciences, and startup funds from the Department of Biomedical Informatics at the University at Buffalo.

### Author’s contributions

ZF, WM, JS, and RS all conceived this project. ZF created the software for interaction scoring and matrix generation, performed analysis of the matrices, and drafted the manuscript. WM created the benchmarking and analysis software. JS provided hypergeometric analysis of the benchmarking data. RS provided mentorship guidance and helped with manuscript development and editing.

## Acknowledgements

We thank the Center for Computational Research (CCR) at the University at Buffalo for providing additional computational resources and other current and former members of the Samudrala group for helpful discussions.

## References

[1] Joseph A DiMasi. New drug development in the united states from 1963 to 1999. Clinical Pharmacology & Therapeutics, 69(5):286–296, 2001.

[2] Joseph A DiMasi, Ronald W Hansen, and Henry G Grabowski. The price of innovation: new estimates of drug development costs. Journal of health economics, 22(2):151–185, 2003.

[3] Alexander Schuhmacher, Oliver Gassmann, and Markus Hinder. Changing r&d models in research-based pharmaceutical companies. Journal of translational medicine, 14(1):105, 2016.

[4] Steven M Paul, Daniel S Mytelka, Christopher T Dunwiddie, Charles C Persinger, Bernard H Munos, Stacy R Lindborg, and Aaron L Schacht. How to improver&d productivity: the pharmaceutical industry’s grand challenge. Nature reviews Drug discovery, 9(3):203, 2010.

[5] Michael S Kinch, Austin Haynesworth, Sarah L Kinch, and Denton Hoyer. An overview of fda-approved new molecular entities: 1827–2013. Drug discovery today, 19(8):1033–1039, 2014.

[6] V Joachim Haupt, Simone Daminelli, and Michael Schroeder. Drug promiscuity in pdb: protein binding site similarity is key. PLoS one, 8(6):e65894, 2013.

[7] Mark Minie, Gaurav Chopra, Geetika Sethi, Jeremy Horst, George White, Ambrish Roy, Kaushik Hatti, and Ram Samudrala. Cando and the infinite drug discovery frontier. Drug discovery today, 19(9):1353–1363, 2014.

[8] Simon K Mencher and Long G Wang. Promiscuous drugs compared to selective drugs (promiscuity can be a virtue). BMC clinical pharmacology, 5(1):3, 2005.

[9] Ismail Kola and John Landis. Can the pharmaceutical industry reduce attrition rates? Nature reviews Drug discovery, 3(8):711, 2004.

[10] Andrew L Hopkins, Jonathan S Mason, and John P Overington. Can we rationally design promiscuous drugs? Current opinion in structural biology, 16(1):127–136, 2006.

[11] William Mangione and Ram Samudrala. Identifying protein features responsible for improved drug repurposing accuracies using the cando platform: Implications for drug design. Preprints, 2018.

[12] Sean Ekins and Antony J Williams. Finding promiscuous old drugs for new uses. Pharmaceutical research, 28(8):1785–1791, 2011.

[13] Yicheng Mei and Baowei Yang. Rational application of drug promiscuity in medicinal chemistry. Future medicinal chemistry, 10(15):1835–1851, 2018.

[14] Ekachai Jenwitheesuk and Ram Samudrala. Identification of potential multitarget antimalarial drugs. JAMA, 294(12):1487–1491, 2005.

[15] Ekachai Jenwitheesuk, Jeremy A Horst, Kasey L Rivas, Wesley C Van Voorhis, and Ram Samudrala. Novel paradigms for drug discovery: computational multitarget screening. Trends in pharmacological sciences, 29(2): 62–71, 2008.

[16] David Cavalla. Predictive methods in drug repurposing: gold mine or just a bigger haystack? Drug discovery today, 18(11-12):523–532, 2013.

[17] Justin Lamb, Emily D Crawford, David Peck, Joshua W Modell, Irene C Blat, Matthew J Wrobel, Jim Lerner, Jean-Philippe Brunet, Aravind Subramanian, Kenneth N Ross, et al. The connectivity map: using gene-expression signatures to connect small molecules, genes, and disease. science, 313(5795):1929–1935, 2006.

[18] Michael J Keiser, Bryan L Roth, Blaine N Armbruster, Paul Ernsberger, John J Irwin, and Brian K Shoichet. Relating protein pharmacology by ligand chemistry. Nature biotechnology, 25(2):197, 2007.

[19] Michael J Keiser, Vincent Setola, John J Irwin, Christian Laggner, Atheir I Abbas, Sandra J Hufeisen, Niels H Jensen, Michael B Kuijer, Roberto C Matos, Thuy B Tran, et al. Predicting new molecular targets for known drugs. Nature, 462(7270):175, 2009.

[20] Eugen Lounkine, Michael J Keiser, Steven Whitebread, Dmitri Mikhailov, Jacques Hamon, Jeremy L Jenkins, Paul Lavan, Eckhard Weber, Allison K Doak, Serge Côté, et al. Large-scale prediction and testing of drug activity on side-effect targets. Nature, 486(7403):361, 2012.

[21] Zheng Zhao, Che Martin, Raymond Fan, Philip E Bourne, and Lei Xie. Drug repurposing to target ebola virus replication and virulence using structural systems pharmacology. BMC bioinformatics, 17(1):90, 2016.

[22] Naiem T Issa, Jordan Kruger, Henri Wathieu, Rajarajan Raja, Stephen W Byers, and Sivanesan Dakshanamurthy. Druggenex-net: a novel computational platform for systems pharmacology and gene expression-based drug repurposing. BMC bioinformatics, 17(1):202, 2016.

[23] Annie P Chiang and Atul J Butte. Systematic evaluation of drug–disease relationships to identify leads for novel drug uses. Clinical Pharmacology & Therapeutics, 86(5):507–510, 2009.

[24] Assaf Gottlieb, Gideon Y Stein, Eytan Ruppin, and Roded Sharan. Predict: a method for inferring novel drug indications with application to personalized medicine. Molecular systems biology, 7(1):496, 2011.

[25] Bin Chen, Ying Ding, and David J Wild. Assessing drug target association using semantic linked data. PLoS computational biology, 8(7):e1002574, 2012.

[26] Rong Xu and QuanQiu Wang. Large-scale extraction of accurate drug-disease treatment pairs from biomedical literature for drug repurposing. BMC bioinformatics, 14(1):181, 2013.

[27] Ruifeng Liu, Narender Singh, Gregory J Tawa, Anders Wallqvist, and Jaques Reifman. Exploiting large-scale drug-protein interaction information for computational drug repurposing. BMC bioinformatics, 15(1):210, 2014.

[28] Halil Bisgin, Zhichao Liu, Reagan Kelly, Hong Fang, Xiaowei Xu, and Weida Tong. Investigating drug repositioning opportunities in fda drug labels through topic modeling. BMC bioinformatics, 13(15):S6, Sep 2012. doi: 10.1186/1471-2105-13-S15-S6.

[29] Allan Peter Davis, Cynthia Grondin Murphy, Robin Johnson, Jean M Lay, Kelley Lennon-Hopkins, Cynthia Saraceni-Richards, Daniela Sciaky, Benjamin L King, Michael C Rosenstein, Thomas C Wiegers, et al. The comparative toxicogenomics database: update 2013. Nucleic acids research, 41(D1):D1104–D1114, 2012.

[30] Brady Bernard and Ram Samudrala. A generalized knowledge-based discriminatory function for biomolecular interactions. Proteins: Structure, Function, and Bioinformatics, 76(1):115–128, 2009.

[31] H.M. Berman, J. Westbrook, Z. Feng, G. Gilliland, T.N. Bhat, H. Weissig, I.N. Shindyalov, and P.E. Bourne. The protein data bank. Nucleic Acids Research, 28:235–242, 2000.

[32] Chengxin Zhang, Peter L. Freddolino, and Yang Zhang. Cofactor: improved protein function prediction by combining structure, sequence and protein–protein interaction information. Nucleic Acids Research, 45(W1): W291–W299, 2017. doi: 10.1093/nar/gkx366.

[33] Ambrish Roy, Jianyi Yang, and Yang Zhang. Cofactor: an accurate comparative algorithm for structure–based protein function annotation. Nucleic Acids Research, 40(W1):W471–W477, 2012. doi: 10.1093/nar/gks372.

[34] Ambrish Roy and Yang Zhang. Recognizing protein–ligand binding sites by global structural alignment and local geometry refinement. Structure, 20(6):987–997, 2012. doi: 10.1016/j.str.2012.03.009.

[35] Noel M. O’Boyle, Michael Banck, Craig A. James, Chris Morley, Tim Vandermeersch, and Geoffrey R. Hutchison. Open babel: an open chemical toolbox. Journal of Cheminformatics, 3(1):33, Oct 2011. doi: 10.1186/17582946-3-33.

[36] Geetika Sethi, Gaurav Chopra, and Ram Samudrala. Multiscale modelling of relationships between protein classes and drug behavior across all diseases using the cando platform. Mini reviews in medicinal chemistry, 15(8):705–717, 2015.

[37] Gaurav Chopra and Ram Samudrala. Exploring polypharmacology in drug discovery and repurposing using the cando platform. Current pharmaceutical design, 22(21):3109–3123, 2016.

[38] Jeremy A Horst, Adrian Laurenzi, Brady Bernard, and Ram Samudrala. Computational multitarget drug discovery. In Polypharmacology in Drug Discovery, chapter 13, pages 236–302. Hoboken, NJ: John Wiley and Sons Publishing Co, 2012. doi: 10.1002/9781118098141.ch13.

[39] James Schuler and Ram Samudrala. Fingerprinting cando: Increased accuracy with structure and ligand based shotgun drug repurposing. In prep., 2018.

[40] Craig Knox, Vivian Law, Timothy Jewison, Philip Liu, Son Ly, Alex Frolkis, Allison Pon, Kelly Banco, Christine Mak, Vanessa Neveu, Yannick Djoumbou, Roman Eisner, An Chi Guo, and David S. Wishart. Drugbank 3.0: a comprehensive resource for ‘omics’ research on drugs. Nucleic Acids Research, 39(suppl_1):D1035–D1041, 2011. doi: 10.1093/nar/gkq1126.

[41] Ruili Huang, Noel Southall, Yuhong Wang, Adam Yasgar, Paul Shinn, Ajit Jadhav, Dac-Trung Nguyen, and Christopher P. Austin. The ncgc pharmaceutical collection: A comprehensive resource of clinically approved drugs enabling repurposing and chemical genomics. Science Translational Medicine, 3(80):80ps16–80ps16, 2011. doi: 10.1126/scitranslmed.3001862.

[42] Qingliang Li, Tiejun Cheng, Yanli Wang, and Stephen H. Bryant. Pubchem as a public resource for drug discovery. Drug Discovery Today, 15(23): 1052–1057, 2010. doi: 10.1016/j.drudis.2010.10.003.

[43] F Garcia-Cordoba, JM Garcia-Santos, G Díaz González, A Garcia-Geronimo, F Zambudio Muñoz, F Hernández Peñalver, and L Baño Aledo Del. Decrease of unnecessary chest x-rays in intensive care unit: application of a combined cycle of quality improvement. Medicina intensiva, 32(2):71–77, 2008.

[44] Wolf Dietrich Ihlenfeldt, Yoshimasa Takahashi, Hidetsugu Abe, and Shin-ichi Sasaki. Computation and management of chemical properties in cactvs: An extensible networked approach toward modularity and compatibility. Journal of chemical information and computer sciences, 34(1):109–116, 1994.

[45] Xemistry chemoinformatics, 2012. URL http://www.xemistry.com.

[46] Jianyi Yang, Renxiang Yan, Ambrish Roy, Dong Xu, Jonathan Poisson, and Yang Zhang. The i-tasser suite: protein structure and function prediction. Nature methods, 12(1):7, 2015. doi: 10.1038/nmeth.3213.

[47] Jianyi Yang, Ambrish Roy, and Yang Zhang. Biolip: a semi-manually curated database for biologically relevant ligand–protein interactions. Nucleic acids research, 41(D1):D1096–D1103, 2012.

[48] Paul CD Hawkins, A Geoffrey Skillman, and Anthony Nicholls. Comparison of shape-matching and docking as virtual screening tools. Journal of medicinal chemistry, 50(1):74–82, 2007.

